# New insights into TNFα/PTP1B and PPARγ pathway through RNF213- a link between inflammation, obesity, insulin resistance and Moyamoya disease

**DOI:** 10.1101/2020.07.07.192153

**Authors:** Priyanka Sarkar, Kavitha Thirumurugan

## Abstract

Diabetic patients are always at a higher risk of ischemic diseases like coronary artery diseases. One such ischemic carotid artery disease is Moyamoya. Moyamoya disease (MMD) has been associated with diabetes Type-I and II and the causality was unclear. RNF213 is the major susceptible gene for MMD. To understand the association between diabetes mellitus and MMD we chose the major players from both the anomalies, insulin and RNF213. But before establishing a role of RNF213 in insulin regulating pathway we had to understand the involvement of RNF213 within different biological systems. For this we have adopted a preliminary computational approach to understand the prominent interactions of RNF213. Our first objective was to construct an interactome for RNF213. We have analyzed several curated databases and adapted a list of RNF213 interacting partners to develop its interactome. Then to understand the involvement of this interactome in biological functions we have analyzed major biological pathways, biological processes and prominent clusters related to this interactome through computational approach. Then to develop a pathway that might give clue for RNF213 involvement in insulin regulatory pathway we have validated the intercluster and intracluster predictions and identified a regulatory pathway for RNF213. RNF213 interactome was observed to be involved in adaptive immunity with 4 major clusters; one of the cluster involved TNFα. Immune system involves several pathways, and therefore at this point we have chosen an event-based strategy to obtain an explicit target. Immunity is mediated by many pro-inflammatory cytokines like TNFα. TNFα-mediated inflammation, obesity and insulin resistance are associated. Therefore we chose to explore the role of RNF213 in TNFα-mediated inflammation in macrophages and inflammation-mediated insulin-resistance in adipocytes. We have observed an enhancement of RNF213 gene expression by LPS mediated pro-inflammatory stimuli and suppression by PPARγ-mediated anti-inflammatory, insulin sensitizing stimuli in macrophages. A more significant response was observed in adipocytes as well. Administration of the pro-inflammatory cytokine TNFα was able to impede the reduction in RNF213 expression during adipogenesis and this effect was observed to be mediated by PTP1B. Inactivation of PTP1B abolished RNF213 expression which in turn enhanced the adipogenesis process through enhanced PPARγ. Constitutive expression of RNF213 suppressed the adipocyte differentiation by the inhibition of PPARγ. We could show the expression of RNF213 has been regulated by TNFα/PTP1B pathway and PPARγ. The constitutive expression of RNF213 during adipogenesis appears to be an adipostatic measure that obese patients acquire to inhibit further adipogenesis. This is verified *in silico* by analyzing the gene expression data obtained from Gene Expression Omnibus database, which showed a higher expression of RNF213 in adipose tissue samples of obese people. Overall this study gives new insights in the TNFα-mediated pathway in adipogenesis and suggests a role of RNF213 in adipogenesis via this pathway.

## Introduction

Diabetic patients have been marked at a higher risk of coronary artery diseases lead by ischemic injury (Howangyin & Silvestre, 2014). This has made Diabetes mellitus a leading cause for stroke and microvasculature impairments in brain (Ergul, Kelly-Cobbs, Abdalla, & Fagan, 2012). Diabetes has been enormously linked to cerebrovascular diseases (Dalal & Parab, 2002; Zhou, Zhang, & Lu, 2014). Moyamoya disease is an ischemic cerebrovascular disease of carotid arteries. RNF213 (Ring Finger Protein 213) the founder susceptible gene for MMD has been extensively studied to elucidate its role in the pathogenesis of Moyamoya disease (Fujimura et al., 2014; Kamada et al., 2011; Kim, 2016; Shoemaker et al., 2015). MMD is characterized by sprouting of vessels at the base of the brain and stenosis of internal carotid artery caused by hyperplasia of smooth muscle cells present in the intima of carotid arteries. This is sometimes accompanied by lipid accumulation in the proliferating intima which ultimately leads to occlusion due to reduction in the lumen space of carotid arteries (J. Suzuki & Takaku, 1969; Yamauchi et al., 2000). This is quite similar to the condition observed in Type 2 diabetes complications leading to stroke (Zhou et al., 2014). Further MMD has been associated to type 2 Diabetes mellitus in some reports, through their clinical investigations (S. Suzuki et al., 2011). Study by Hatasu Kobayashi, suggested the involvement of RNF213 in type I Diabetes mellitus. They showed that ablation of RNF213 retarded the progression of diabetes in Akita mice (H. Kobayashi et al., 2013). Akita mice are model for type I Diabetes mellitus with a mutation in Ins2 (Pre-proinsulin 2).

RNF213 is an E3 ubiquitin ligase with AAA^+^ ATPase domain and a RING domain to perform the ligase activity (Morito et al., 2014). Though most of the previous studies had focused on its physiological and clinical aspects, few independent studies suggested potential regulatory mechanism for RNF213. Study by Scholz suggested RSPO3 (R-spondin3) as a co-regulatory gene for RNF213 (Scholz et al., 2016). Another study by Kazuhiro Ohkubo suggested that RNF213 is transcriptionally activated by the synergistic effect of TNFα and IFNγ in endothelial cells, and PKR and PI3K-AKT pathways act as upstream regulators for these cytokines. Also they revealed the involvement of RNF213 in inflammation through detailed analysis of curated datasets (Ohkubo et al., 2015). It is still not known whether these cytokines directly regulate the transcription of RNF213 or indirectly through some downstream regulators. Further, RNF213 protein was reported to be a substrate for PTP1B (Banh et al., 2016). PTP1B is a negative regulator of insulin (Nieto-Vazquez et al., 2007). TNFα is also known to cause insulin resistance (Lorenzo et al., 2008). It also acts as an anti-adipogenic factor in a way through altering PTP1B (D. D. Song et al., 2013). Also, these cytokine-mediated pro-inflammatory molecules are secreted by activated macrophages.

When a host is invaded by a pathogen, dendritic cells are the first to get triggered, followed by macrophages. Activated macrophages and dendritic cells act as effector phase molecule for the adaptive immunity by engulfing, processing and presenting the antigens on its surface to T_H_ cells and activates inflammation (Cronkite & Strutt, 2018; Janeway, P, M, & Al., 2001; N. F. and K. Kobayashi, 2005). Inflammation has been extensively studied in relation to obesity. Though obesity is stated as a low grade inflammation, pro-inflammatory cytokines are known to act as negative regulators for adipocyte differentiation.

All these studies gave valuable insights about the regulatory mechanism that might cue the involvement of RNF213. But a detailed analysis of the plausible interactome for RNF213 has not been performed.

Therefore, we have adopted an *in silico* approach to predict an interactome for RNF213. Gene co-regulatory and gene ontology studies have always been valued for predicting the functional attributes of a gene. Several tools are available online to predict accurate hits which can further be screened and validated and we applied this methodology as a base for our study. Based on these findings we have designed the study to validate some of our *in silico* predictions and explored new insights into an already existing anti-adipogenic insulin regulatory pathway.

## Results

### *In silico* analysis

The predicted interactome of RNF213 (Figure 1) was observed to be involved in several biological systems (Figure2a) but mainly involved in immunity and cytokine driven bioprocesses (Figure 2b) among which MHC Class1 antigen processing and presentation in immune system was the major hit (Figure 2c). There were four major gene clusters (Figure 3a) observed to be functioning within the interactome. One of the clusters belongs to RNF213 and it had 12 members including RNF213. Among the observed list of proteins, DTX3L, TRIM21 and HERC6 were co-regulated with RNF213 (Supplementary data 3). Each cluster belongs to members having similar function within a biological system. Among these 4 clusters, other 2 clusters belong to members involved in inflammation and host defense immune responses. One cluster included NOTCH1 (Fazio & Ricciardiello, 2016; Toshihiro Ito, Judith M. Connett, Steven L. Kunkel, 2012) and the other cluster belong to TNFα and PTP1B (G. J. Song et al., 2016; Zabolotny et al., 2008). The fourth cluster represented the ATP synthase members. Macrophages were selected as *in vitro* model to validate the *in silico* predictions. Macrophages are the key effectors and modulator cells of immune system (Martinez & Gordon, 2014). Raw 264.7 murine macrophages were chosen because they are activated on encountering pathogens similar to dendritic cells. Macrophages engulf these pathogens and digest the antigen into smaller peptides which are presented to CD8^+^ T cells on MHC class I molecules (Cronkite & Strutt, 2018). Macrophages are also known to secrete inflammatory molecules and trigger inflammation directly (Janeway et al., 2001; N. F. and K. Kobayashi, 2005). Therefore, macrophages were chosen as an efficient model for intercluster and intracluster validation.

**Figure 1.**
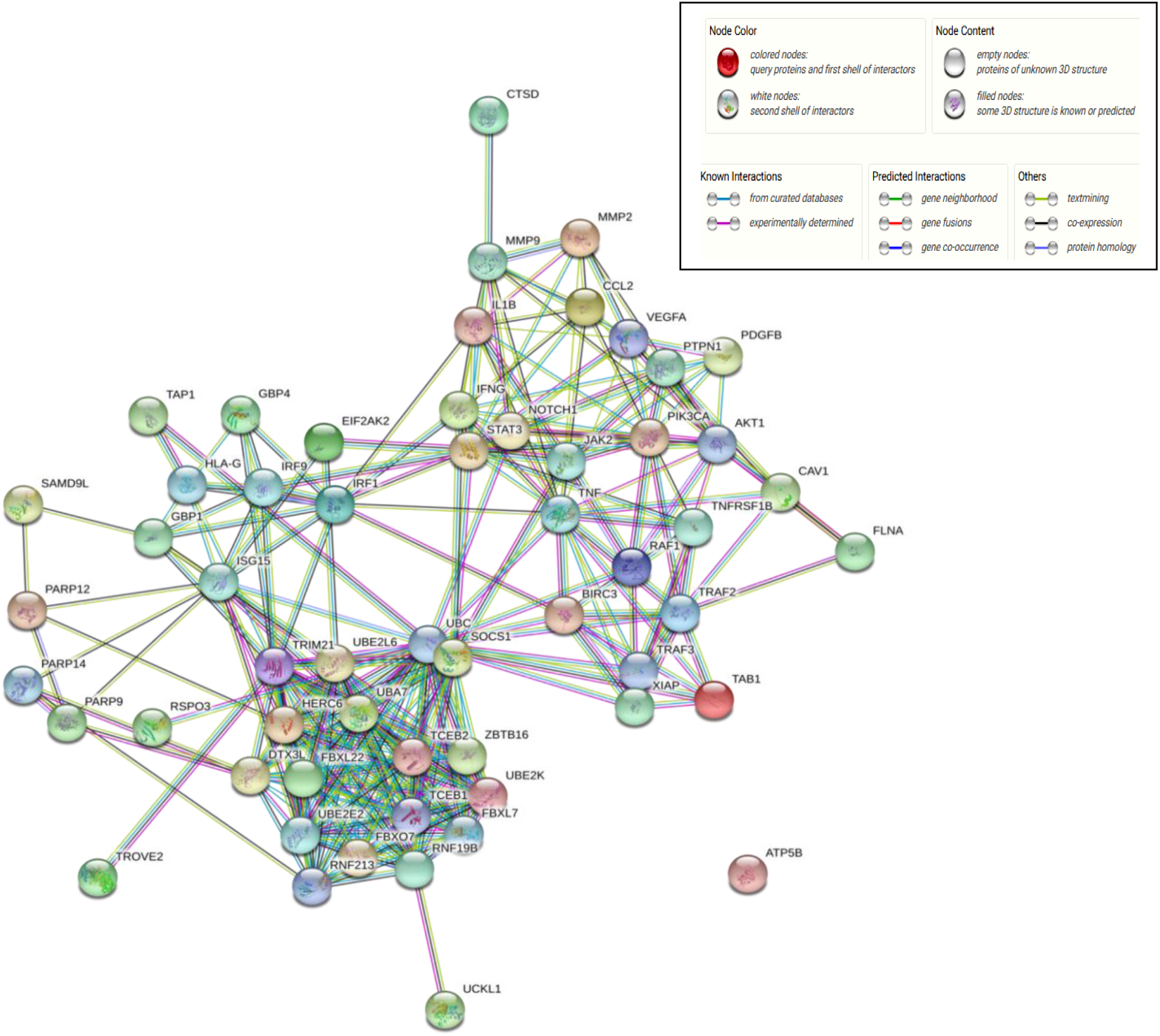
Predicted interactome of RNF213. The candidates of the interactome were grouped together in STRING database to visualize their interactions

**Figure 2.**
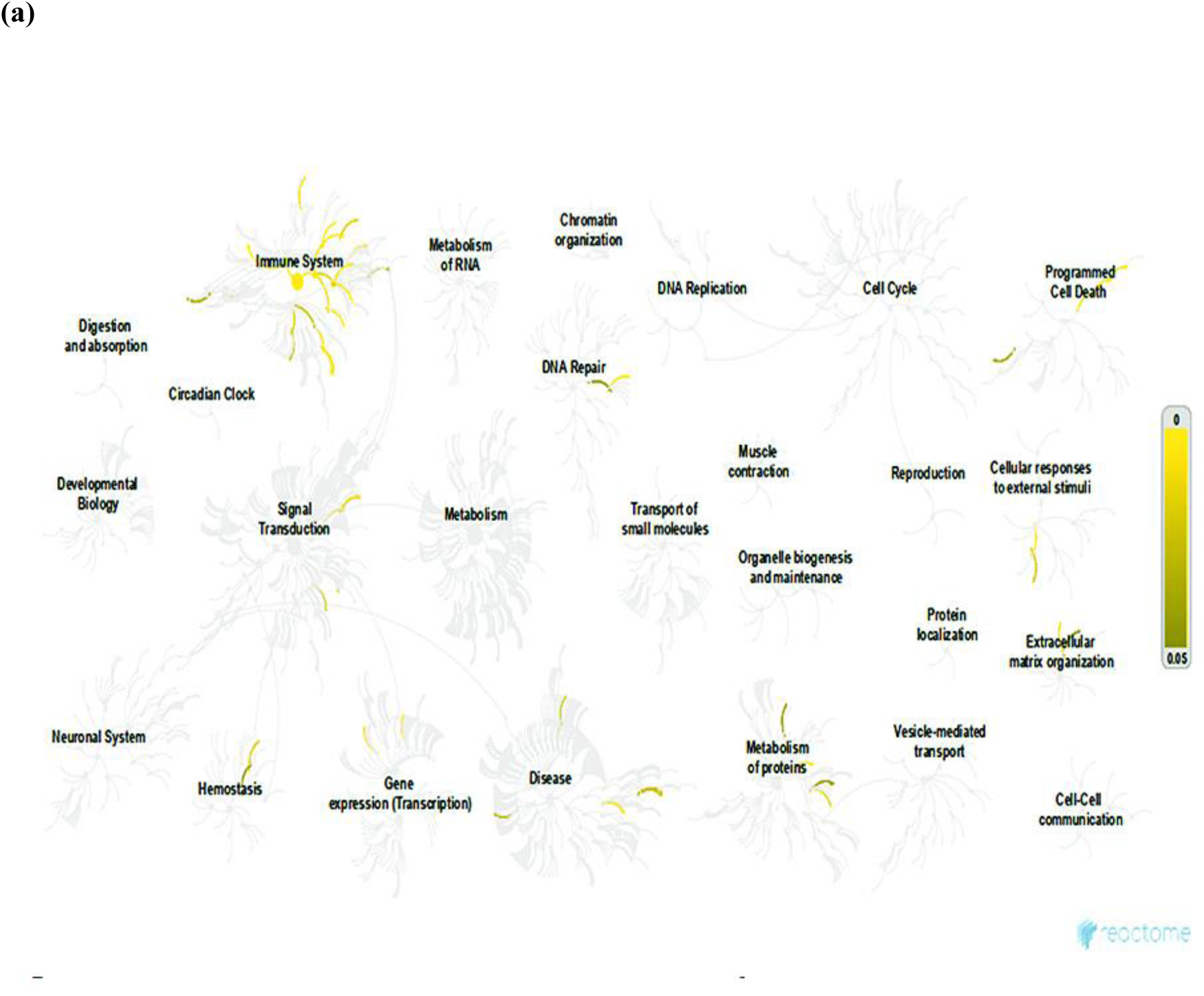
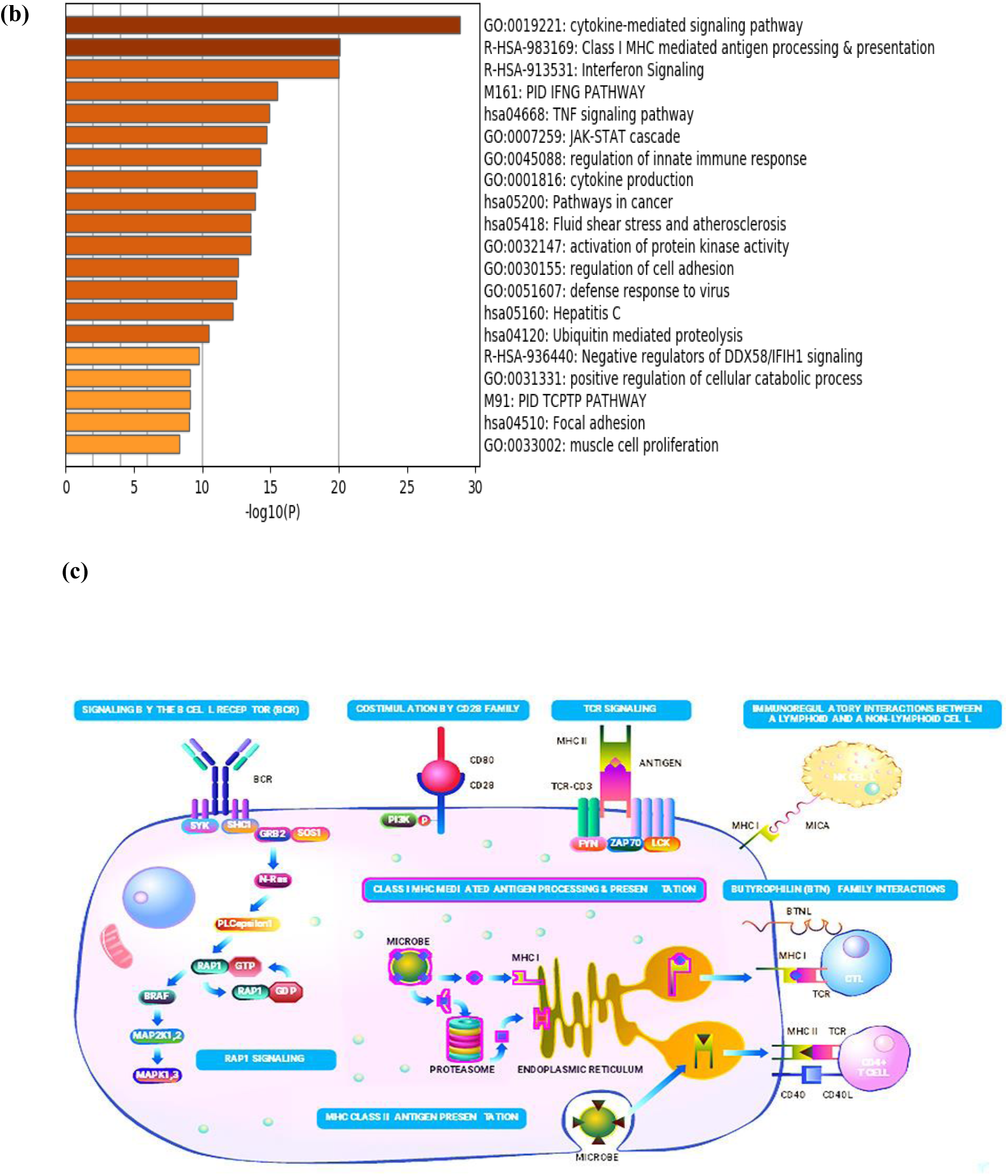
(a) Involvement of predicted interactome in several biological systems. (b) Enrichment process showing the predominance of interactome in immunity and inflammation. (c) Involvement of RNF213 in MHC class I antigen processing. Dynamic areas of RNF213 are encased in pink.

**Figure 3.**
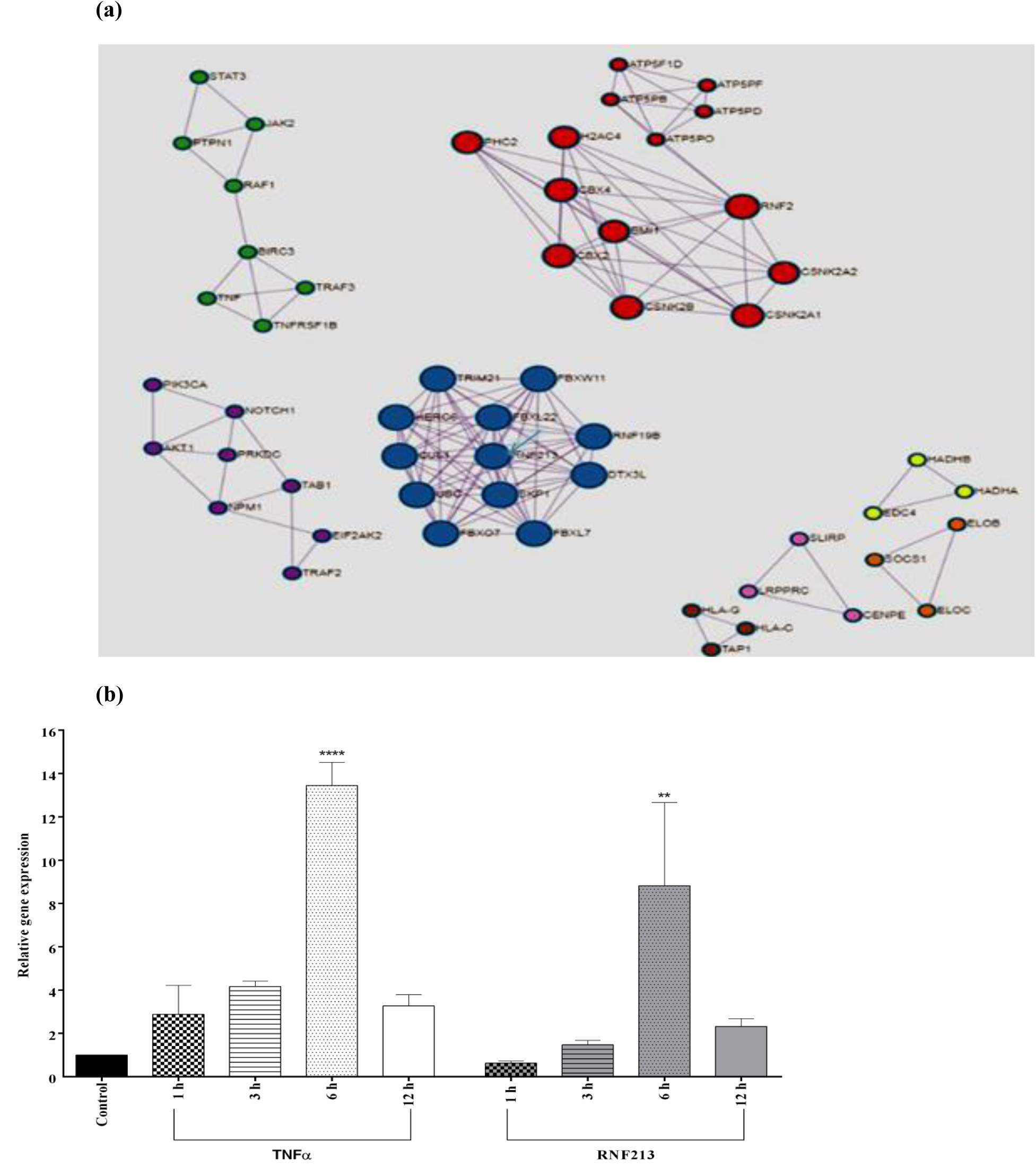
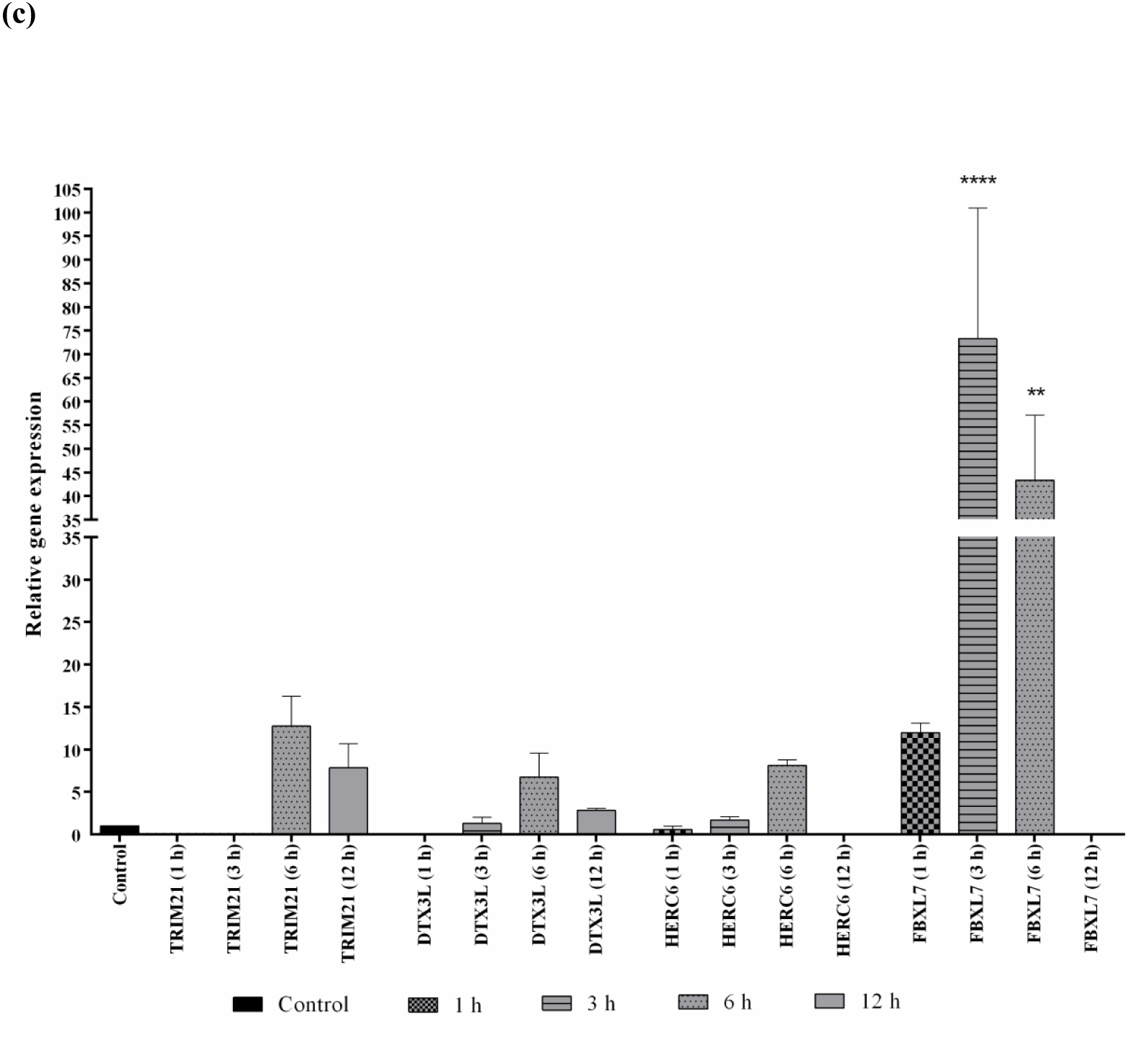
(a) There were four major gene clusters observed to be functioning within the interactome of RNF213. Each color represents a different biological function. (b) Treatment of Raw 264.7 cells with LPS induced the expression of TNFα and RNF213 and enhanced expression was noticed after six hours of activation (b) Expression profile of co-regulated members of RNF213 cluster (TRIM21, DTX3L, HERC6) and non-co-regulated member FBXL7 after LPS treatment.

### Intercluster and intracluster validation

Raw 264.7 cells were stimulated with LPS. LPS induces classical activation of macrophages (Martinez & Gordon, 2014) by enhancing the secretion of pro-inflammatory cytokine, TNFα (Reis et al., 2012; Soromou et al., 2012). When these cells were treated with LPS, it induced the expression of RNF213 at transcriptional level. Expression of TNFα and RNF213 was pronounced after six hours of activation with LPS (Figure3**b**). Along with this, the co-regulated members of RNF213 cluster were also analyzed to check whether they too show similar expression profile on LPS stimulation. Interestingly, HECT and RLD domain containing E3 ubiquitin protein ligase family member 6(HERC6), Tripartite motif-containing protein 21 (TRIM21), Deltex E3 Ubiquitin Ligase 3L (DTX3L) displayed a similar expression profile (Figure3**c**) to that of RNF213 and TNFα (Fig.3**b**). Thus members of RNF213 cluster were properly grouped as they were regulated in a similar fashion by the inflammatory stimulus, thereby validating the intra-cluster grouping. In contrast to the co-regulated genes, F-box/LRR-repeat protein 7 (FBXL7) though being a member of the same cluster had some variations with respect to the pattern of RNF213 expression pattern (Figure 3**b**). Similarity between TNFα expression pattern from the other cluster and the expression pattern of members of RNF213 cluster, indicates an intercluster interaction. At this stage we have concluded that TNFα individually might also be able to regulate RNF213.

TNFα is stated as an interlinking node between insulin resistance, obesity and inflammation. It mediates Wnt and inflammation signaling to prevent adipocyte differentiation (Gustafson & Smith, 2006) in 3T3-L1 by suppressing adipogenic genes (Ruan, Hacohen, Golub, Parijs, & Lodish, 2002) and also by impeding the reduction of PTP1B (D.D. Song et al., 2013).

### RNF213 Expression in Adipocytes

Therefore to link inflammation and adipogenesis we have attempted to evaluate the expression of RNF213 in adipocytes. We observed RNF213 was expressed well during the first 2 days of adipocyte differentiation and on the 8th day (Figure 4a). There was an inclined suppression of RNF213 expression from 4^th^ day onwards up to 6^th^ day. This is the time when the preadipocyte differentiate into mature adipocytes. In parallel to the regulation of RNF213 we also profiled PTP1B expression that was observed to be similar to that of RNF213 expression during adipocyte differentiation (Figure 4b). This suggested an involvement of major adipogenic regulators like PPARγ and CEBPα in RNF213 regulation. Therefore, we have evaluated the effect of relation between RNF213, PPARγ and CEBPα expression pattern. CEBPα was not considered for further analysis because it was not synchronized with the expression profile of RNF213 (data not shown). Therefore, we have evaluated the effect of PPARγ on RNF213.

**Figure 4.**
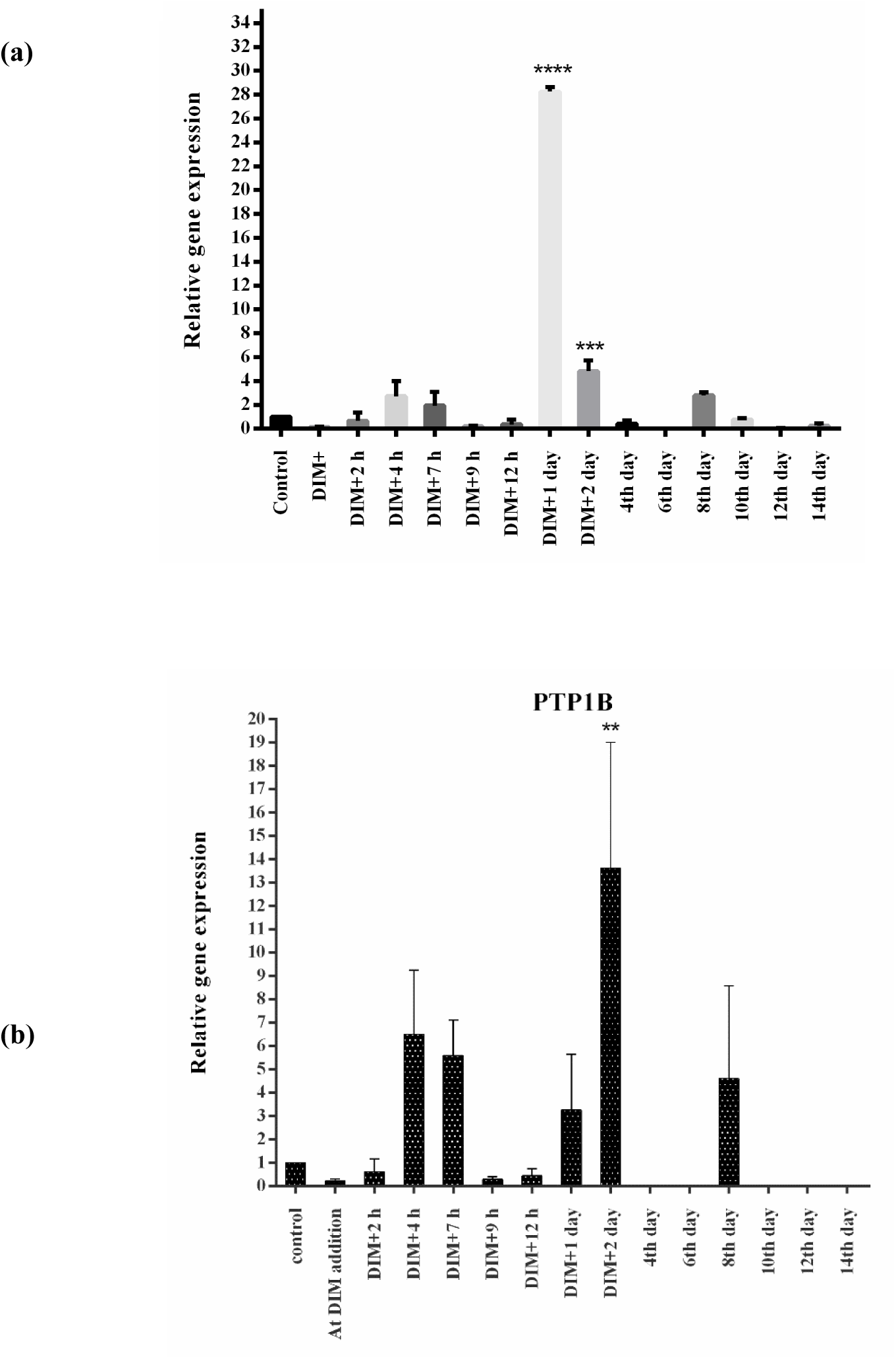
(a)Expression of RNF213 at mRNA level during adipocyte differentiation. RNF213 expressed well during the first 2 days of adipocyte differentiation and on the 8th day. (b) Expression of PTP1Bat mRNA level during adipocyte differentiation. PTP1B expression pattern was similar to that of RNF213.

### Suppression of RNF213 by PPARγ agonist

PPARγ is a master regulator of adipogenesis and it is activated by thiazolidinediones. Here we have administered pioglitazone. Pioglitazone is not only an activator of PPARγ but it also acts as an anti-inflammatory molecule by suppressing TNFα expression both at protein and mRNA level by activating PPARγ and inactivating NFκB (Ao et al., 2010). It also suppresses the expression of IFNγ in a PPARγ dependent manner(Cunard et al., 2019). Further it decreases the insulin resistance (Kemnitz et al., 1994). Therefore, we have evaluated the effect of pioglitazone treatment in macrophages as well as in adipocytes.

RNF213 was induced by inflammation and slightly suppressed by PPARγ dependent anti-inflammation in macrophages (Figure 5a). Pioglitazone acted as an anti-inflammatory molecule by completely suppressing the expression of TNFα (Figure 5a). Further, there was a significant reduction in the mRNA expression of RNF213 in pioglitazone treated 3T3-L1 adipocytes (Figure 5b). These results indicate insulin sensitivity and anti-inflammation might negatively regulate RNF213 gene expression. We did not use PPARγ inhibitor or PPARγ - RNAi at this stage because we wanted to evaluate the RNF213 expression pattern throughout the adipogenesis process. But inhibiting PPARγ will block adipogenesis.

**Figure 5.**
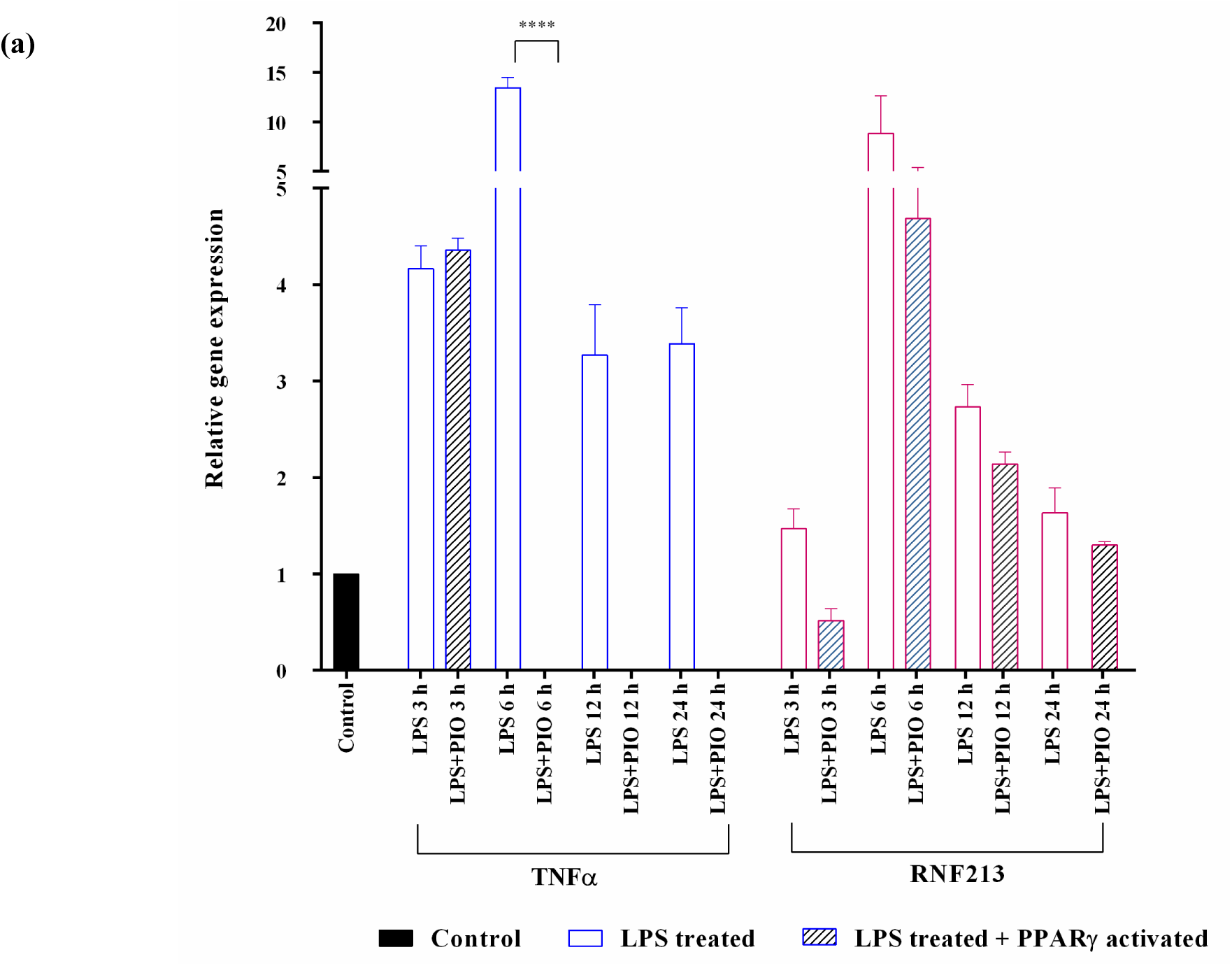
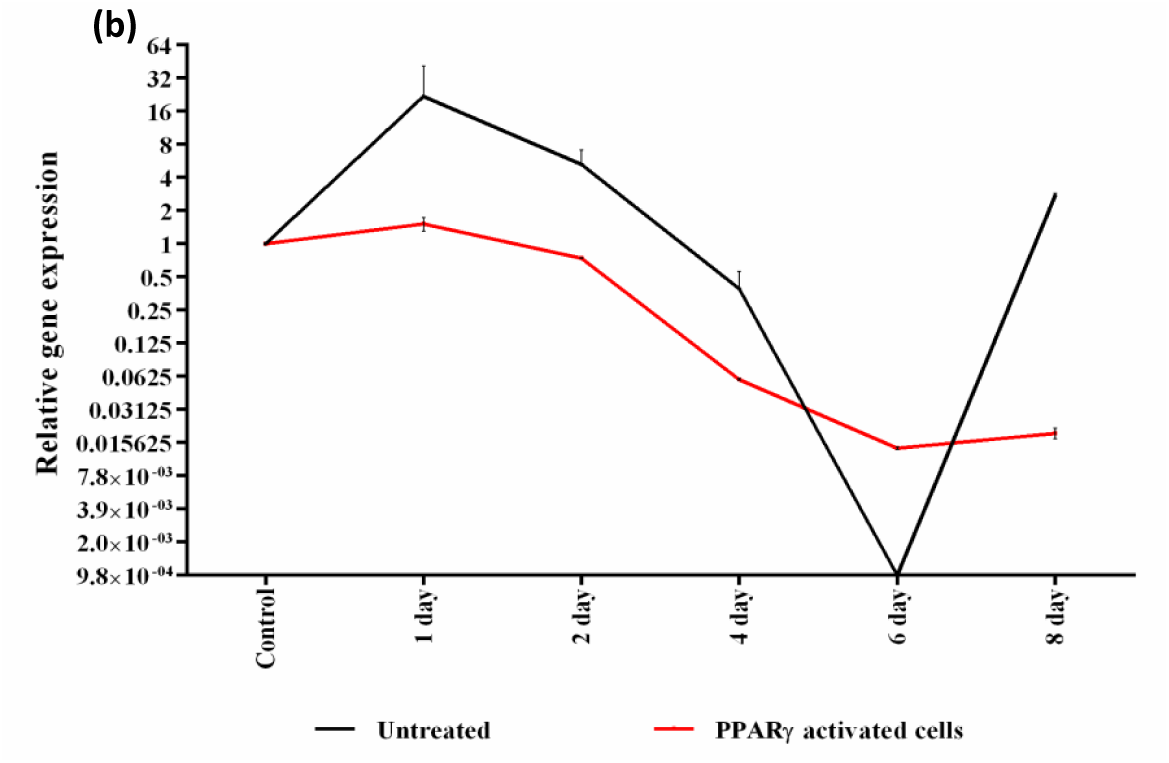
(a) Effect of pioglitazone on mRNA expression of TNFα and RNF213. LPS stimulated Raw 264.7 cells treated with pioglitazone showed significantly reduced expression of TNFα and slightly reduced expression of RNF213. (b) Expression of RNF213 in 3T3-L1 adipocyte cells treated with pioglitazone. Pioglitazone activates PPARγ which in turn reduced the RNF213 expression.

### Effect of TNFα on RNF213 expression

We have attempted to evaluate the effect of pro-inflammatory, negative regulator of insulin, TNFα on RNF213 expression. For this, 3T3-L1 pre-adipocytes were treated with TNFα at an inflammatory dose causing adipostatic effect (Gustafson & Smith, 2006). The treatment of TNFα impeded the reduction of RNF213 mRNA throughout adipogenesis (Figure 6a). We again performed a parallel expression profiling for PTP1B. We observed a similar pattern in PTP1B expression to that of RNF213 expression (Figure 6b). The same trend was seen in the protein expression of RNF213 and PTP1B following TNFα treatment (Figure 7a-e). Immunostained adipocyte cells were observed for RNF213 and PTP1B expression at day 2 and day 5 of differentiation. The cells treated with TNFα expressed RNF213 and PTP1B throughout adipogenesis process (Figure 7a-d).

**Figure 6.**
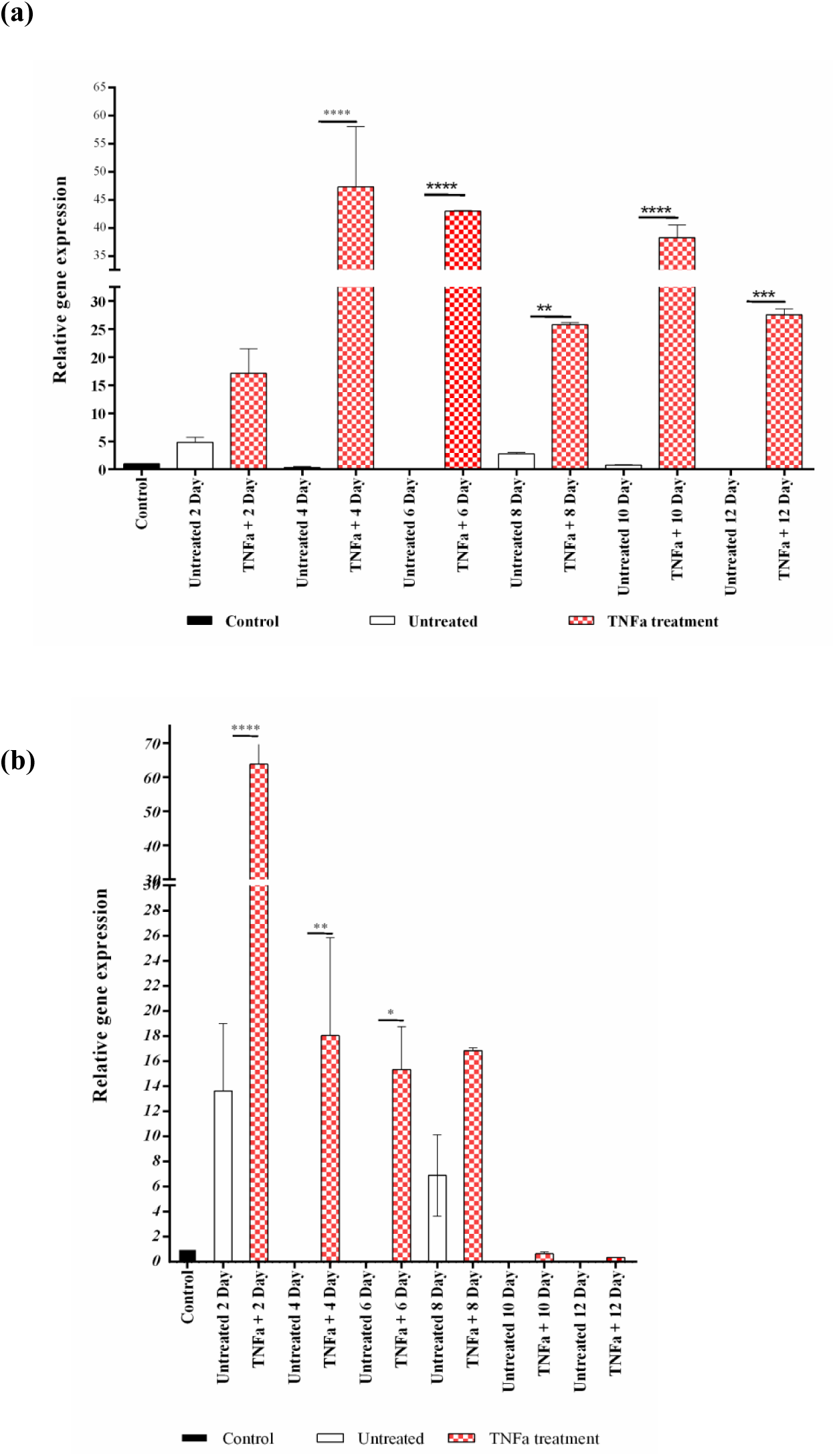
(a) RNF213 expression in 3T3-L1 adipocytes treated with TNFα. Administration of TNFα (1.5 ng/ml) along with the differentiation media causes a constitutive expression of RNF 213 throughout the adipogenesis process up to 12 days (b) PTP1B expression in 3T3-L1 adipocytes treated with TNFα. Similar to RNF213, the expression of PTP1B was high throughout the adipogenesis process up to 8 days.

**Figure 7.**
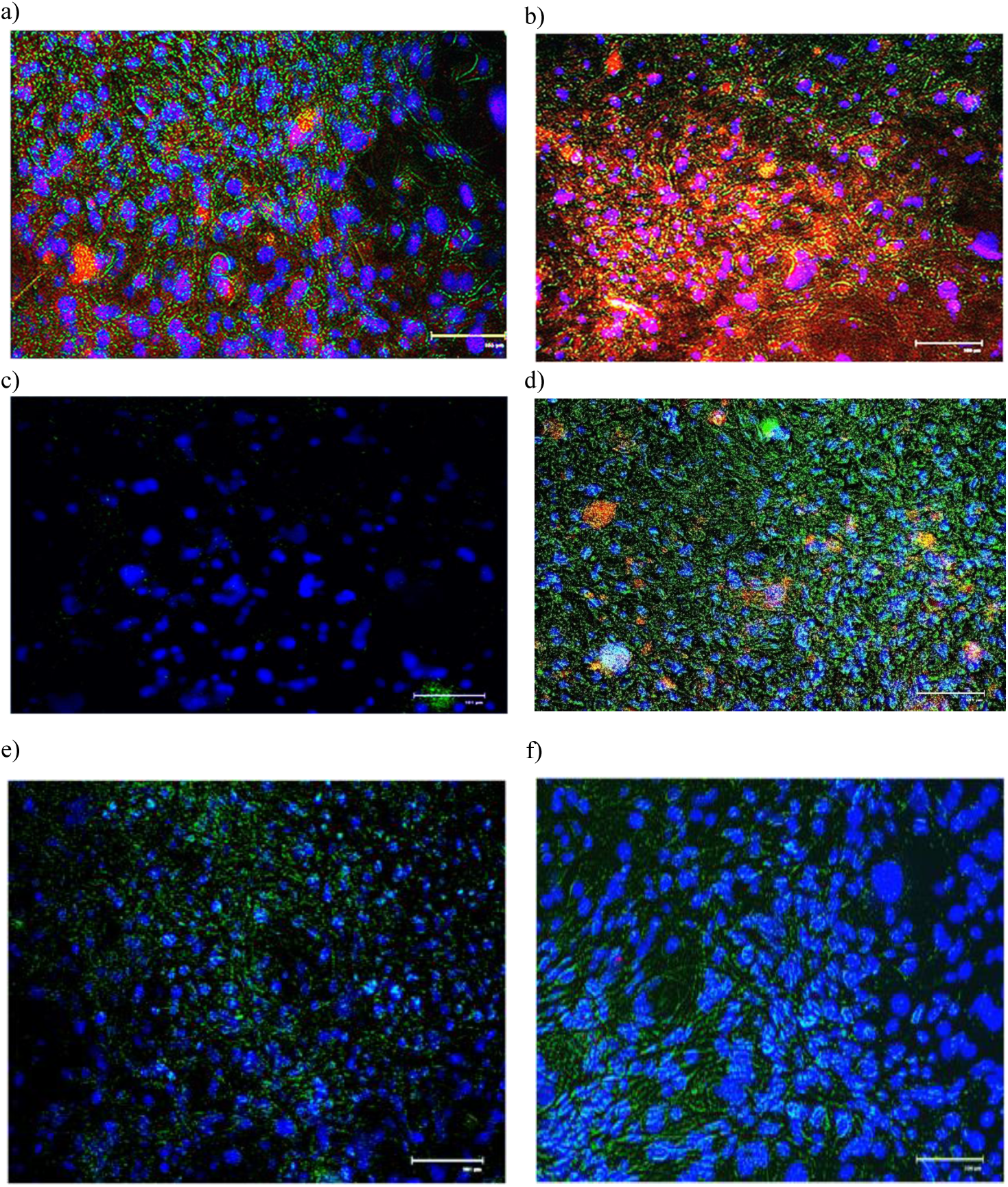
Immunoprecipitation was performed with fluorescently labeled antibodies for invitro protein expression analysis a) RNF213 and PTP1B expression in 3T3-L1 adipocyte cells at day 2 of differentiation. **b)** RNF213 and PTP1B expression in TNFα treated adipocytes at day 2 of differentiation. **c)** RNF213 and PTP1B expression in adipocytes at day 5 of differentiation. **d)** RNF213 and PTP1B expression in TNFα treated adipocytes at day 5 of differentiation. The expression of RNF213 and PTP1B was increased by TNFα administration and stayed constitutive during the adipogenesis process. **e)** RNF213 and PTP1B expression in a TNFα and TCS401 co-treated adipocytes at day 5 of differentiation. **f)** RNF213 and PTP1B expression in adipocyte cells at day 2 of differentiation treated with only TCS401.The effect of TNFα on the constitutive expression of RNF213 was nullified when PTP1B was inactivated. RNF213 was indicated in red colour (Alexa488 tagged antibody), PTP1B was indicated in green colour (Cy3 tagged antibody) and the samples were counterstained with DAPI.

### Effect of PTP1B on RNF213 expression

Further we wanted to investigate the mechanism followed by TNFα. Since PTP1B was reported as one of the downstream partners of TNFα insulin resistance pathway and in our data also it showed similar trend of expression to that of RNF213 expression. We have evaluated the effect of PTP1B on RNF213. For this we have analysed the effect of PTP1B inactivation on RNF213 expression. The administration of sodium orthovanadate (phosphatase inhibitor) at 35 µM suppressed the mRNA expression of RNF213 (Figure8a). Whereas using PTP1B inhibitor TCS401 specifically at 0.29 μM concentration abolished the expression of RNF213 at gene level (Figure 8b) and protein level. This was indicated by *in vitro* protein expression analysis measured with fluorescently labelled antibodies (Figure7f). Further when TNFα treated cells were co-treated with TCS401, it nullified the TNFα mediated enhanced effect on RNF213 expression (Figure 7e). Day 8 adipocytes treated with TNFα show reduced adipogenesis indicated by less number of Oil Red O stained lipid droplets (Figure 9a-ii) and PPARγ transcript levels (Figure 9b). Same cells with TCS401 co-treatment show enhanced number of lipid droplets suggesting the role of PTP1B on adipogenesis (Figure 9a-iii).

**Figure 8.**
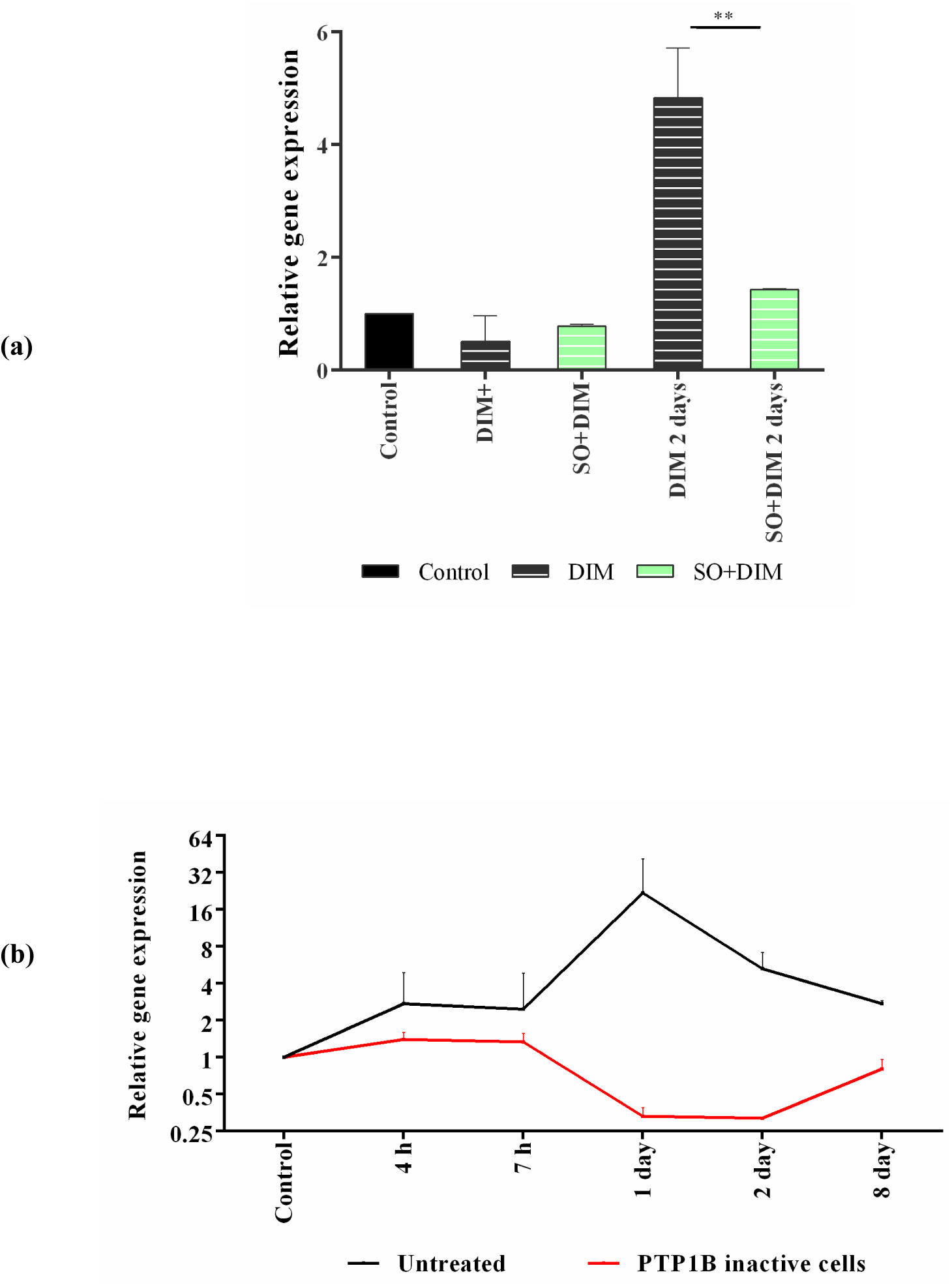
(a) Sodium orthovanadate treatment of adipocytes. Sodium orthovanadate at 35 µM reduced the expression of RNF213 at day 2 of differentiation (b) RNF213 expression in TCS401 treated cells. PTP1B specific inhibitor TCS401 abolished the expression of RNF213 as it inactivated PTP1B at 0.29 μM concentration.

**Figure 9.**
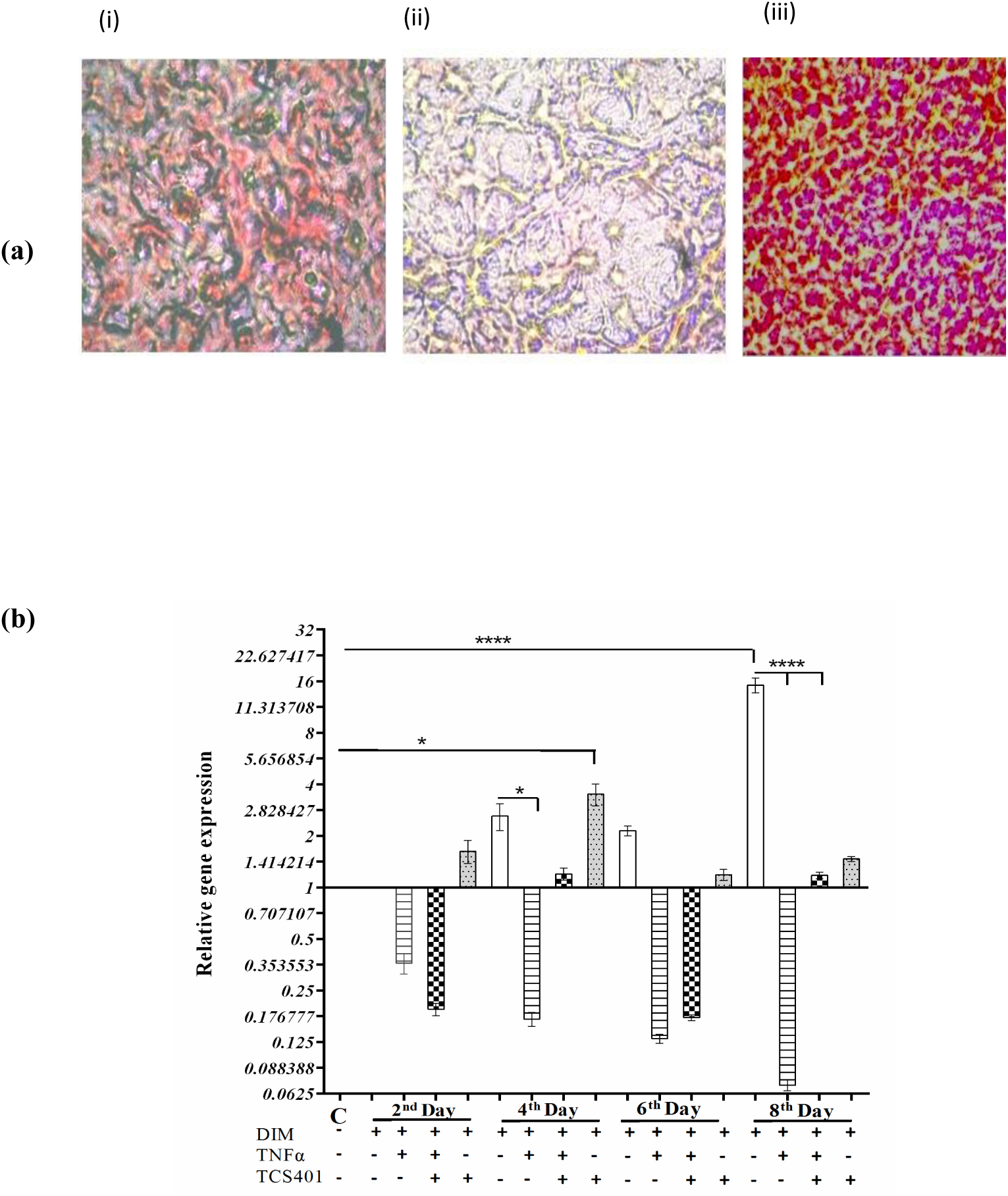
(a) Oil Red O staining of adipocytes at day 8 of differentiation.. (**i)** Normal adipogenesis process. (**ii)** TNFα treated cells display less number of lipid droplets indicating reduced adipogenesis. (**iii)** TNFα treated cells with TCS401 co-treatment show enhanced number of lipid droplets suggesting the role of PTP1B on adipogenesis. **b)** Gene expression pattern of PPARγ in adipocytes treated with DIM (differentiation induction media), TNFα, and TCS401. PPARγ expression was gradually increasing and it reached the maximum at day 8 in the cells treated with DIM. PPARγ expression was very low in the cells treated with TNFα. This effect was reversed in cells treated with TCS401.

This indicated that TNFα can also regulate RNF213 and it is mediated through PTP1B. This data showed a complete regulatory dependence of RNF213 on PTP1B.

### Effect of positive regulators of RNF213 (TNFα and PTP1B) on PPARγ

Since RNF213 expression was observed to be suppressed by activation we wanted to analyze the effects of positive regulators of RNF213 on PPARγ, the negative regulator of RNF213.

First we have evaluated the gene expression pattern of PPARγ in normal differentiating cells and then compared it with the TNFα treated differentiating cells and PTP1B inhibited differentiating cells. Gene expression of PPARγ was high in adipocytes treated with DIM (differentiation induction media) particularly at day 8 (Figure 9b). PPARγ expression was very low in the cells treated with TNFα. This effect was reversed in cells treated with PTP1B specific inhibitor TCS401 (Figure 9b), the downstream partner of TNFα. This shows that administration of TNFα suppresses PPARγ in the presence of PTP1B. TNFα affects several adipogenic molecules through different pathways. Therefore PPARγ could be acting downstream to PTP1B or it might have its own regulatory pathway to modulate RNF213 expression.

The inactivation of PTP1B individually also increased adipogonesis as indicated by the increased number of lipid droplets (Figure 10) and PPARγ expression levels (Figure 9b). Therefore PTP1B might be regulating RNF213 through PPARγ and further analysis is required to evaluate this mechanism.

**Figure 10.**
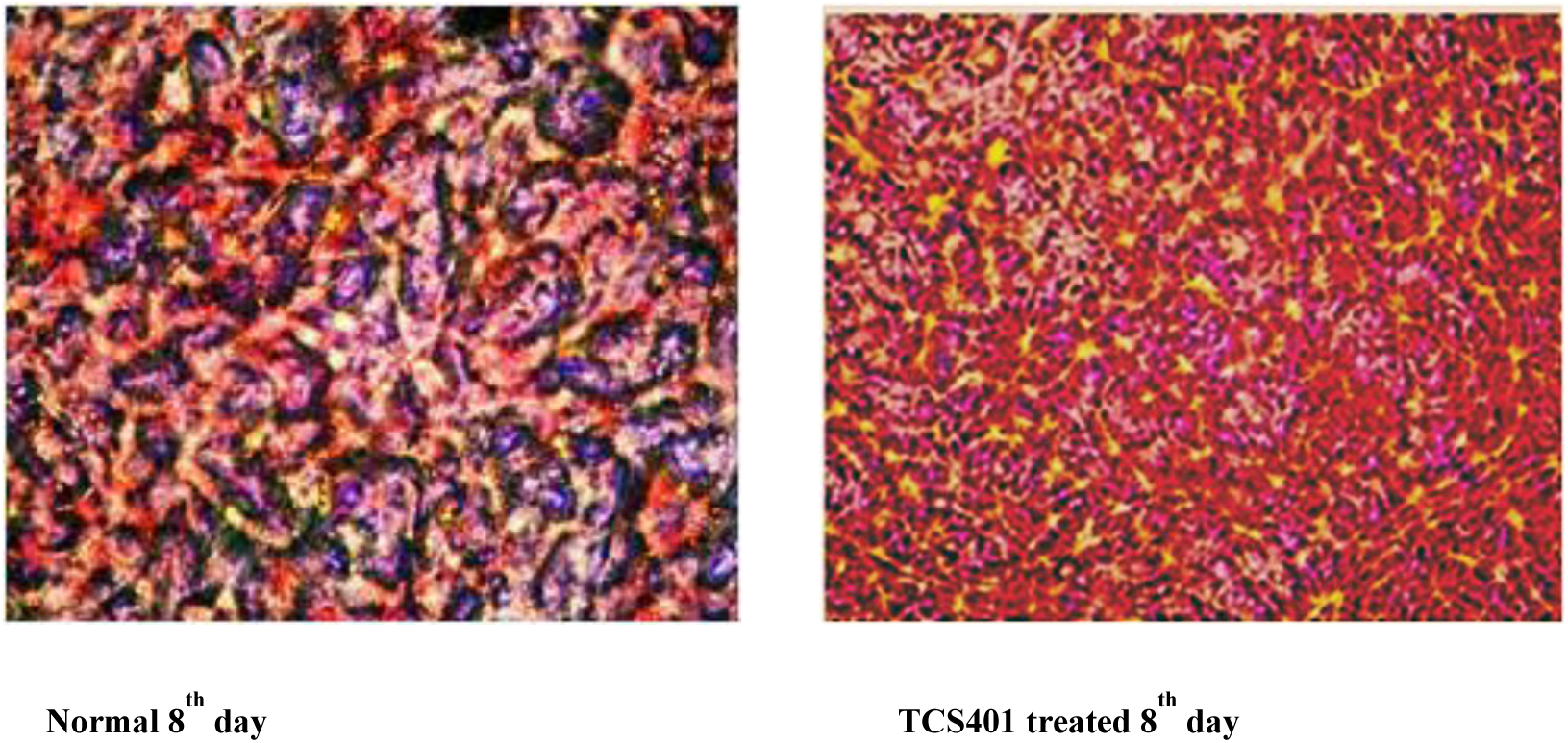
Oil Red O staining was performed on the 8^th^ day for TCS401 treated differentiated adipocytes. Adipogenesis was increased when PTP1B was inactivated. This was confirmed by the increased levels of PPARγ expression as depicted in Figure 8b

### Gene expression analysis from Microarray database

Our data suggest the involvement of RNF213 in adipocyte differentiation. This is in accordance to the gene expression data curated from Gene Expression Omnibus through GEO2R tool, where we can observe a 4.5 fold increase in RNF213 expression in non-diabetic obese PIMA individuals (Supplementary data 4). This was the only curated normalized dataset available that significantly recognized RNF213.

Most of the other datasets had variable probes for RNF213 with unreliable Padj values. Therefore we chose this dataset for our evaluation. Obesity is said to be a low grade/chronic inflammation which leads to insulin resistance (Choi & Cohen, 2017). An increase in the gene expression of RNF213 during obesity suggests its role in obesity related insulin resistance and predicts its likely protective effect in obese patients by reducing adipogenesis.

### Perspectives and Conclusion

It is decisive to mark the interacting partners for the gene to know the exact role and the regulatory mechanism of a gene. RNF213 is observed to be present across many uncurated datasets, making it relevant to list the plausible interactors. By curating such datasets we have observed that RNF213 shows an increased gene expression during obese conditions. RNF213 has been reported to be induced by inflammatory stimuli (Ohkubo et al., 2015) and its ablation improves glucose tolerance (H. Kobayashi et al., 2013). These reports are in accordance with our study showing the induction of RNF213 expression by TNFα/PTP1B inflammatory pathway leading towards adipostatic effects. This pathway is known to be involved in insulin resistance in adipocytes (Lorenzo et al., 2008). Our data also shows the continuous expression of RNF213 downstream to PTP1B suppresses adipogenesis. But its ablation by PTP1B inactivation increases adipogenesis. Further there is a cyclic pattern of RNF213 expression suggesting the existence of a feedback inhibition mechanism to regulate RNF213. In this study, we had predicted a whole curated interactome for RNF213 which is partly validated. This prediction highlighted the emerging role of RNF213 in inflammation and inflammation mediated anti-adipogenesis. Also we had speculated that TNFα/PTP1B pathway positively regulates RNF213 expression and negatively regulates adipogenesis. Further RNF213 knockdown analysis is required to confirm the influential role of RNF213 in TNFα/PTP1B mediated insulin resistance and adipostasis.

From our data it was clear that a reduction in RNF213 expression was required to achieve adipogenesis and this reduction was caused by the activation of PPARγ; indicating PPARγ as an effective regulator of RNF213. RNF213 is expressed in both macrophages as well as adipocytes. Both these cell types are related to inflammation. TNFα looks like the common regulator of RNF213 via PTP1B but PPARγ appears to be more effective in suppressing RNF213 in adipocytes suggesting a link between adipogenesis, insulin resistance, inflammation and MMD via TNFα, PTP1B, PPARγ and RNF213. PTP1B might be inactivating PPARγ in order to induce RNF213, and TNFα and PTP1B is known to suppress PPARγ. Further analysis is required to state whether PPARγ modulates RNF213 through this pathway or some other pathway. Overall TNFα/PTP1B insulin-resistant pathway enhances RNF213 expression whereas PPARγ mediated insulin sensitization suppresses its expression. Therefore RNF213 could be another link between obesity, inflammation, insulin resistance and MMD like TNFα.

## Methods

### Interactome prediction

A molecular interactome was predicted using protein interactors and gene interactors. Interacting genes were listed down from UCSC Genome Browser’s gene interaction tool and GeneMania. Protein-protein interactions were based on literature survey, physical interactors, co-expressed partners and functional homology transfers. Physical interactors and co-expressed partners were sorted through GeneMania and STRING. The functional homologs were detected through Genedecks online web tool. These homologous functional partners were sorted based on domain matching and the protein interactors for these partners were listed as homology transfer interactions. All the interactions were sorted based on their probability matching and false discovery rate (FDR). The value 0.01 was considered as a cutoff for FDR. List of these molecules (Supplementary data 1) were used to predict a molecular interactome dataset and it was uploaded on STRING database to develop a visual interacting network with high confidence (0.7). A diagrammatic representation of this approach has been given in the attachments as flow chart for interactome prediction.

### Interactome validation and pathway prediction

The obtained interactome dataset was submitted to METASCAPE to obtain enrichment cluster analysis and functional complexes through EXPRESS Analysis. METASCAPE results were verified through DAVID. Members of the interactome dataset were submitted to REACTOME to confirm the biological systems and pathway analysis. These results were confirmed through KEGG PATHWAY by considering the KEGG ontology terms for these molecules. The major complexes as presented by the use of MCODE algorithm were based on the top non-redundant enriched terms from METASCAPE and the biological system pathway from REACTOME. The members of these complexes were analyzed for their co-regulation through Database of Gene Co-Regulation (dGCR) (Williams, 2015) and validated for their co-regulated expression and ligand stimulated regulatory pathway in cell lines.

### Cell culture

RAW 264.7 cells and 3T3-L1 cells were bought from NCCS (Pune) cell repository, India. RAW 264.7 cells were cultured in DMEM (Himedia) and 10% FBS (Invitrogen, USA) media containing 1% antibiotic-antimycotic solution (100X, Gibco) as described previously (George, Ramasamy, & Sirajudeen, 2019). The 3T3-L1 pre-adipocytes were grown as previously described (Shihabudeen, Roy, James, & Thirumurugan, 2015). Briefly, cells were grown for 2 days post confluence in DMEM (Invitrogen, USA) supplemented with 10% new born calf serum (Invitrogen, USA). Differentiation was then induced by changing the medium to DMEM supplemented with 10% fetal bovine serum, 0.5 mM 3-isobutyl-1-methylxanthine, 1 µM dexamethasone, and 1.2 µM insulin (Sigma-Aldrich, USA). After 48 h, the differentiation medium (referred to as DIM) was replaced with maintenance medium containing DMEM supplemented with 10% fetal bovine serum and 1.2 µM insulin for 48 hours post induction. Thereafter maintenance medium was replaced every 48 h until 14 days. Cells were incubated at 37°C in a 5% CO_2_ environment.

### Treatment and sample collection

Raw 264.7 cells were treated with 1µg/ml of LPS to induce inflammation. These cells were collected at different time points; post induction (1h, 3h, 6h, 12h and 24h) and gene expression for the samples were normalized against samples from un-induced Raw 264.7 cells. PPARγ activation for these cells was done by treating the cells with 10µM concentration of pioglitazone.

Samples for 3T3-L1 cells were collected at different time points after inducing with differentiation media (at hours (h) 0h, 2h, 4h, 7h, 9h, 12h, 24h, 48h, and at days 4, 6, 8, 10, 12, 14). Sodium orthovanadate (Sigma) was administered at a concentration of 35 µM as reported previously along with DIM (Liao & Lane, 1995). TNFα (Sigma) was administered at a concentration of 1.5 ng/ml along with DIM and maintenance media up to 12 days. This concentration was chosen based on the previous reports of TNFα causing inflammatory response in 3T3-L1 cells (Gustafson & Smith, 2006).TCS401, specific inhibitor of PTP1B (Veda scientific) (Iversen et al., 2000) was administered at a concentration of 0.29 µM along with DIM and maintenance media up to 8 days. Gene expression for the samples collected at different time points were normalized against samples from 48 hours pre-induction (referred to as −2 day or control) for 3T3-L1 cells.

### Gene expression

Total RNA was isolated using TRIzol reagent according to manufacturer’s instructions (Invitrogen, USA). The cDNA was synthesized from 1 µg of total RNA using the Prime script cDNA conversion kit (Takara, India). Gene expression was measured using SYBR green dye (Takara, India) in BIORAD CFX96 Touch Real-Time PCR Detection System. GAPDH was used as an endogenous control in the comparative cycle threshold (CT) method. The list of primers from Xcelris is given in the Supplementary data 2.

### Oil Red O staining

Oil Red O staining was performed following a modified protocol previously described (Kraus et al., 2016). Briefly, cells were fixed with 10% formalin for 45 minutes followed by a washing step with 60% isopropanol. Then the cells were air dried and incubated with Oil Red O stain (Sigma) for 30 minutes. Then the cells were washed properly with distilled water and the dried wells were used for imaging. Imaging was done through Olympus Magnus Phase contrast microscope.

### Immunocytochemistry

3T3-L1 adipocyte cells were fixed with 4% PFA (paraformaldehyde) for 15 minutes at room temperature, washed with PBS and permeabilized with 0.3 % of TRITON X-100 for 10 min. After blocking the cells with blocking buffer (0.3% TRITON X-100, 1% BSA) for 1 hour; the cells were incubated for 5 hours at 4°C with specific primary antibody (Alexa488 tagged RNF213 primary antibody and Cy3 tagged PTP1B primary antibody from BIOSS, USA) and then counterstained with DAPI. Samples were collected on day 1, day 2, day 5 and day 7. Imaging was done by using EVOS FLoid imaging station (Thermo Fischer, USA) with 20x fluorite objective and LED light cubes containing hard coated filters (blue, red and green).

### Gene expression analysis from Microarray database

RNF213 differential expression was re-analyzed in human samples through GEO2R. GEO2R is the R-package offered by Gene Expression Omnibus (GEO) to obtain differentially expressed gene list in a given microarray dataset based on the Fold change and AdjP-values. For this study we have used the dataset accession number GSE2508 from GEO database. GSE2508 dataset comprises of RNA samples isolated from the adipocytes of abdominal subcutaneous fat of non-diabetes Pima Indians (Y. H. Lee, S. Nair, E. Rousseau, P. A. Tataranni, C. Bogardus, P. A. Permana, 2006).

### Statistical analysis

Each experiment had a lower limit of n=3 (3 biological replicates with 3 technical replicates taken as average).Some experiments were repeated more number of times to confirm accuracy. All data were presented as mean ± SEM. Column statistics and ANOVA was performed using Graphpad Prism v.06 software package. *P*< 0.05 was considered to be statistically significant. It is presented as ns (*p*> 0.05), *(*p* ≤ 0.05), **(*p* ≤ 0.01), ***(*p* ≤ 0.001) **** (*p* ≤ 0.0001).

## Acknowledgements

This work has been financially supported by Vellore Institute of Technology’s Seed grant and Senior Research Fellowship by ICMR (Indian Council of Medical Research).

## Conflict of interest

The authors declare that they have no conflict of interest.

---

Supplementary data 1. List of interacting molecules to predict a molecular interactome dataset of RNF213

Supplementary data 2.List of forward and reverse primers used in the experiment Supplementary data 3. Co-regulated members of RNF213 cluster

Supplementary data 4. Curated gene expression data of non-diabetic obese PIMA individuals obtained from Gene Expression Omnibus

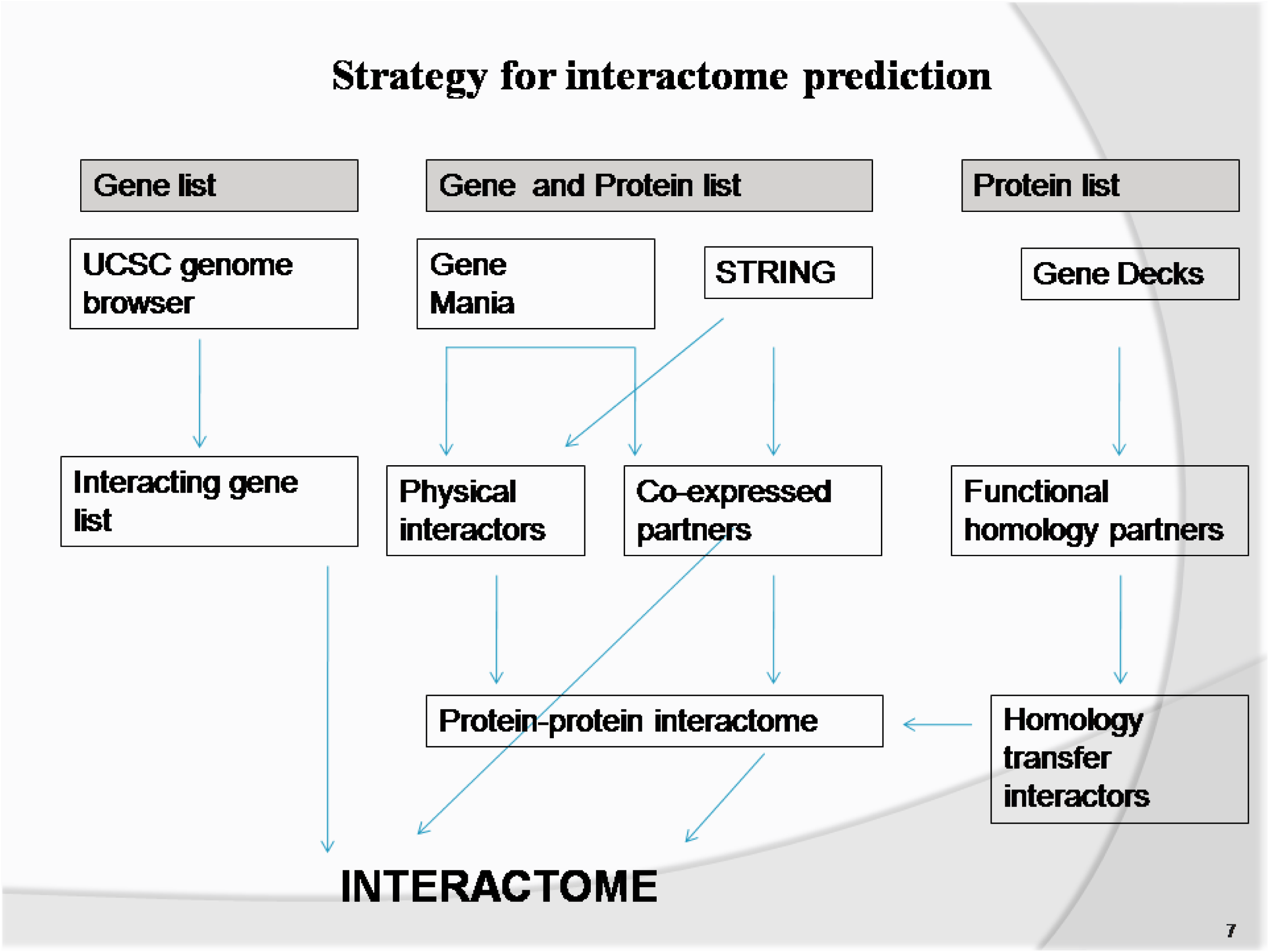

